# Molecular Dynamics Simulations of the Interaction of Mouse and *Torpedo* Acetylcholinesterase with Covalent Inhibitors Explain Their Differential Reactivity: Implications for Drug Design

**DOI:** 10.1101/532754

**Authors:** Nellore Bhanu Chandar, Irena Efremenko, Israel Silman, Jan M.L. Martin, Joel L. Sussman

**Affiliations:** Dept. of Organic Chemistry, Weizmann Institute of Science, 7610001 Reḥovot, Israel.; Dept. of Neurobiology, Weizmann Institute of Science, 7610001 Reḥovot, Israel; Dept. of Structural Biology and Israel Center for Structural Proteomics, Weizmann Institute of Science, 7610001 Reḥovot, Israel.

## Abstract

Although the three-dimensional structures of mouse and *Torpedo californica* acetylcholinesterase are very similar, their responses to the covalent sulfonylating agents benzenesulfonyl fluoride and phenylmethylsulfonyl fluoride are qualitatively different. Both agents inhibit the mouse enzyme effectively by covalent modification of its active-site serine. In contrast, whereas the *Torpedo* enzyme is effectively inhibited by benzenesulfonyl fluoride, it is completely resistant to phenylmethylsulfonyl fluoride. A bottleneck midway down the active-site gorge in both enzymes restricts access of ligands to the active site at the bottom of the gorge. Molecular dynamics simulations revealed that the mouse enzyme is substantially more flexible than the *Torpedo* enzyme, suggesting that enhanced ‘breathing motions’ of the mouse enzyme relative to the *Torpedo* enzyme might explain why phenylmethylsulfonyl fluoride can reach the active site in mouse acetylcholinesterase, but not in the *Torpedo* enzyme. Accordingly, we performed docking of the two sulfonylating agents to the two enzymes, followed by molecular dynamics simulations. Whereas benzenesulfonyl fluoride closely approached the active-site serine in both mouse and *Torpedo* acetylcholinesterase in such simulations, phenylmethylsulfonyl fluoride was able to approach the active-site serine of mouse acetylcholinesterase - but remained trapped above the bottleneck in the case of the *Torpedo* enzyme. Our studies demonstrate that reliance on docking tools in drug design can produce misleading information. Docking studies should, therefore, also be complemented by molecular dynamics simulations in selection of lead compounds.

**Author summary:** Enzymes are protein molecules that catalyze chemical reactions in living organisms, and are essential for their physiological functions. Proteins have well defined three-dimensional structures, but display flexibility; it is believed that this flexibility, known as their dynamics, plays a role in their function. Here we studied the neuronal enzyme acetylcholinesterase, which breaks down the neurotransmitter, acetylcholine. The active site of this enzyme is deeply buried, and accessed by a narrow gorge. A particular inhibitor, phenylmethylsulfonyl fluoride, is known to inhibit mouse acetylcholinesterase, but not that of the electric fish, *Torpedo*, even though their structures are very similar. A theoretical technique called molecular dynamics (MD) shows that the mouse enzyme is more flexible than the *Torped*o enzyme. Furthermore, when the movement of the inhibitor down the gorge towards the active site is simulated using MD, the phenylmethylsulfonyl fluoride can reach the active site in the mouse enzyme, but not in the *Torpedo* enzyme, in which it remains trapped midway down the gorge. Our study emphasizes the importance of taking into account not only structure, but also dynamics, in designing drugs targeted towards proteins.

## 1. Introduction

The principal biological role of acetylcholinesterase (AChE) is termination of transmission at cholinergic synapses by rapid hydrolysis of the neurotransmitter, acetylcholine (ACh) [1, 2].

The crystal structure of *Tc*AChE revealed that, despite the high catalytic activity of AChE, which approaches diffusion control [3], its active site is near the bottom of a long and narrow gorge, >15 Å long, a large part of the surface of which is lined by aromatic residues [4]. Near the mid-point of the gorge is a bottleneck, between two conserved aromatic residues, whose cross-section is smaller than the diameter of the quaternary group of ACh, 6.4 Å. Thus, in *Tc*AChE, the cross-section at the narrowest point of the bottleneck is ~5 Å [4]. In mAChE, the narrowest point of the bottleneck has a cross-section of 2.4 Å [5]. Fig 1 displays the crystal structure of the complex with ACh of the S203A mutant of mAChE. Space-filling representations of two ACh molecules are seen, one lodged above the gorge, and one below it [6]. This representation clearly illustrates that the AChE molecule needs to ‘breathe’ substantially in order for the ACh molecule to pass through the bottleneck to reach the active site.

**Fig 1.**
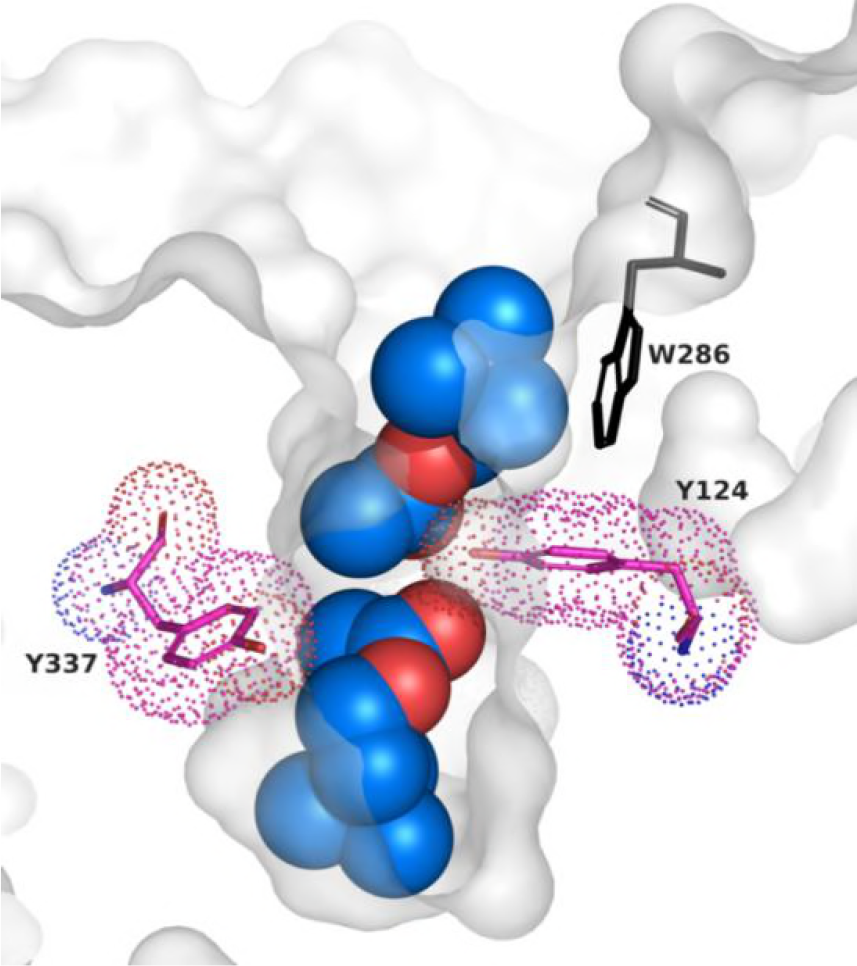
Space-filling representations of two ACh molecules in the active-site gorge of mAChE. The illustration is based on the PDB ID 2HA4 crystal structure [6]. The two Achs are positioned above and below the bottleneck residues, Y337 and Y124, which are displayed as sticks, with magenta dots displaying their full space-filling surface; the surface of the gorge as a whole is displayed in grey.

AChE is the target of a repertoire of inhibitors that act by covalent modification of its active-site serine. These include organophosphate (OP) nerve agents, and both OP and carbamate insecticides [7]. Phenylmethylsulfonyl fluoride (PMSF) is a sulfonylating agent commonly employed as a non-specific inhibitor of serine proteases [8]. It was reported by Fahrney & Gold [9] that *Electrophorus electricus* AChE is resistant to inhibition by PMSF, although it was later reported that it is indeed inhibited, but extremely slowly [10]. It was subsequently shown that mammalian AChEs are very susceptible to inhibition by PMSF [11, 12].

In an earlier study, we reported that AChE from the electric organ of another electric fish, *Torpedo californica* (*Tc*) AChE, is also resistant to PMSF, but is irreversibly inhibited very effectively by its homolog benzenesulfonyl fluoride (BSF) (Scheme 1) [13]. In contrast, we found that mouse AChE (mAChE) is very well inhibited by both PMSF and BSF. These observations are puzzling because the crystal structures of *Tc*AChE [4] and mAChE [14] are very similar (Fig 2), as are the kinetic constants for their extremely rapid action on their natural substrate, ACh, and on its homolog, acetylthiocholine [15, 16]. We subsequently reported that the anti-Alzheimer drug, rivastigmine (Exelon™), carbamylates human AChE >1,600-fold faster than *Tc*AChE [17].

**Scheme 1:**
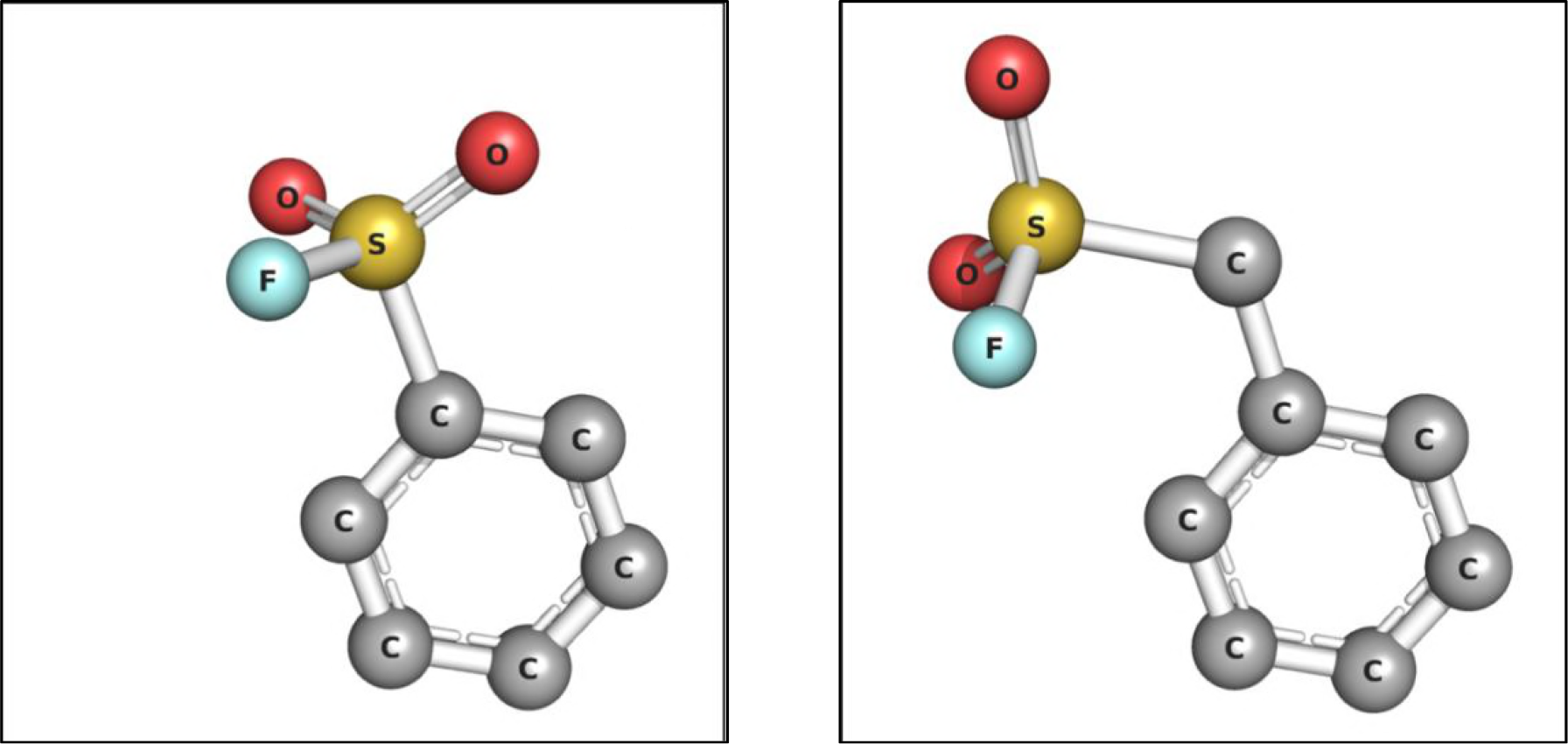
Stick representations of benzenesulfonyl fluoride (BSF, left) and phenylmethylsulfonyl fluoride (PMSF, right) (see S1 Fig in Supplementary Information).

**Fig 2.**
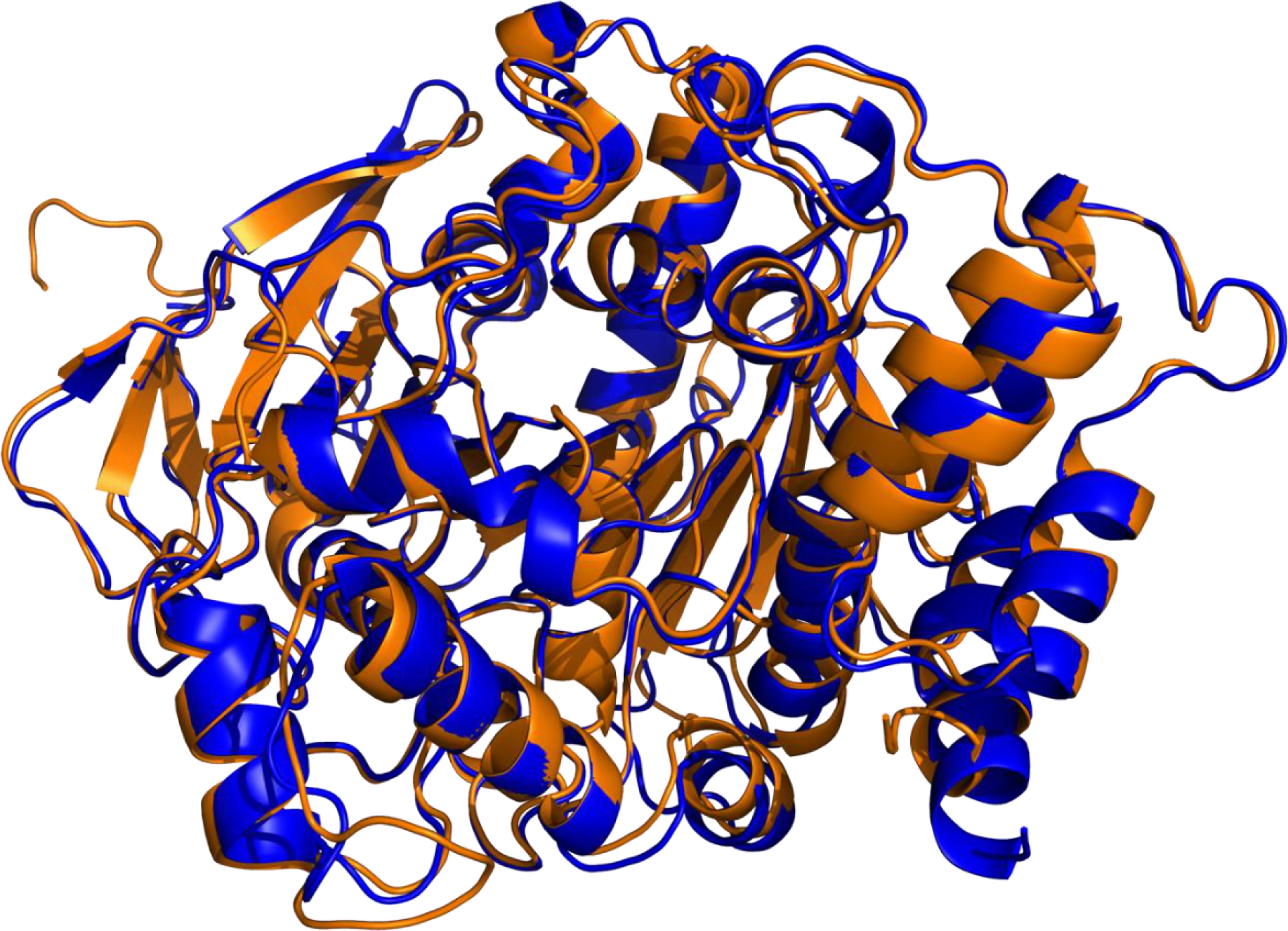
Overlay of cartoon representations of the crystal structures of *Tc*AChE (blue) and mAChE (orange) looking down the active-site gorge.

In the following we present theoretical evidence, using rigid docking in tandem with molecular dynamics (MD) simulations, which shows that differences in flexibility of *Tc*AChE and mammalian AChEs can account for the striking differences experimentally observed in their rates of inhibition by both sulfonylating and carbamylating agents [13, 17, 18]. Thus, reliance on docking data alone in the context of drug design has the potential to produce very misleading results.

## 2. Results

### 2.1. Structural Comparison of *Tc*AChE and mAChE

The overall structures of the two enzymes closely resemble each other, with RMSD values of Cα atoms 0.54 Å for 449 atoms after protein preparation, and 0.54 Å, for 434 atoms, for the RSCB pdb structures. However, careful inspection reveals three clear differences between the *Tc*AChE and mAChE structures:

i. There is a difference in one residue at the bottleneck of the active-site gorge, F330 in *Tc*AChE being replaced by Y337 in mAChE, with the sidechains of these two residues being oriented very differently (Fig 3).
ii. In mAChE there are four residues, P258-P259-G260-G261, which are absent in *Tc*AChE (Fig 4). As a result, a significantly larger loop protrudes in mAChE than in *Tc*AChE (Fig 5).
iii. In addition to the bottleneck residue, four additional residues in the upper portion of the gorge differ in *Tc*AChE and mAChE: E73(T75), Q74(L76), S81(T83), S124(A127).

**Fig 3.**
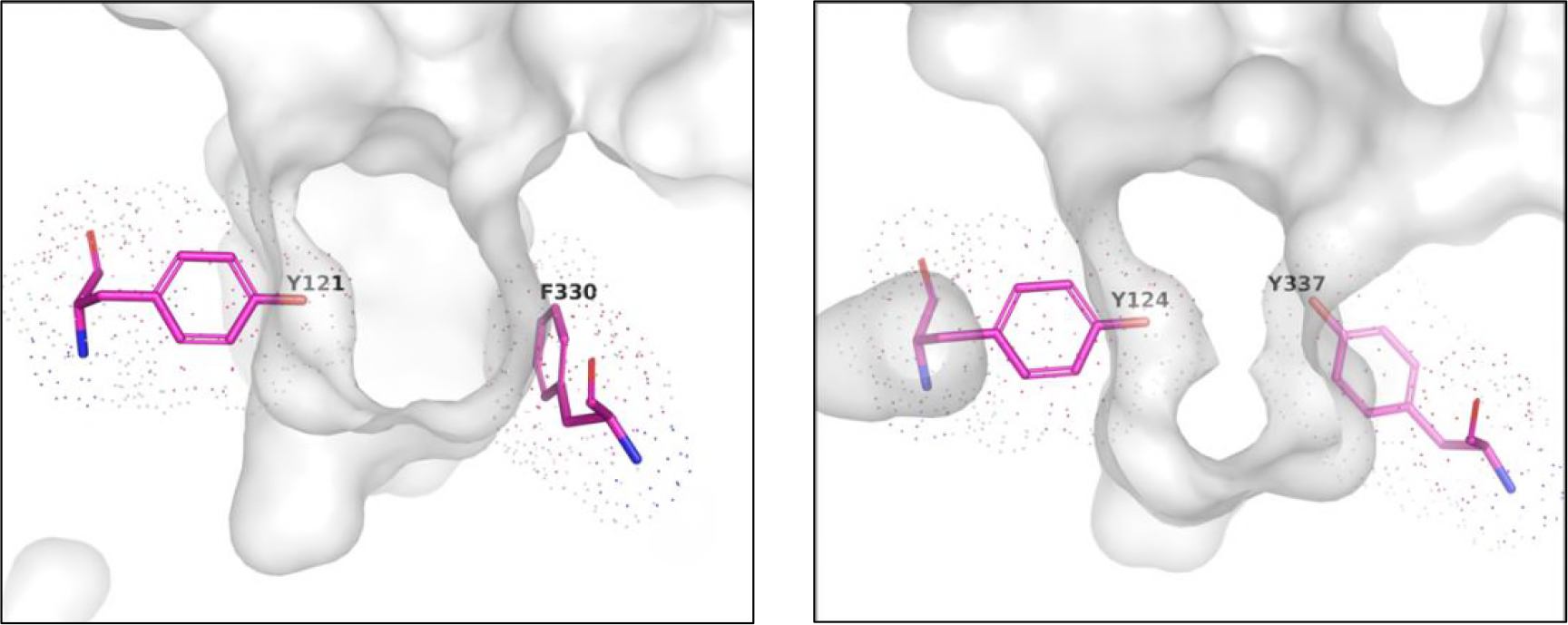
Comparison of the 3D structures of *Tc*AChE and mAChE in the bottleneck region of the active-site gorge after performance of the protein preparation protocol. The two panels show the orientations of the bottleneck residues Y121(Y124)/F330(Y337) of *Tc*AChE (left) and mAChE (right).

**Fig 4.**
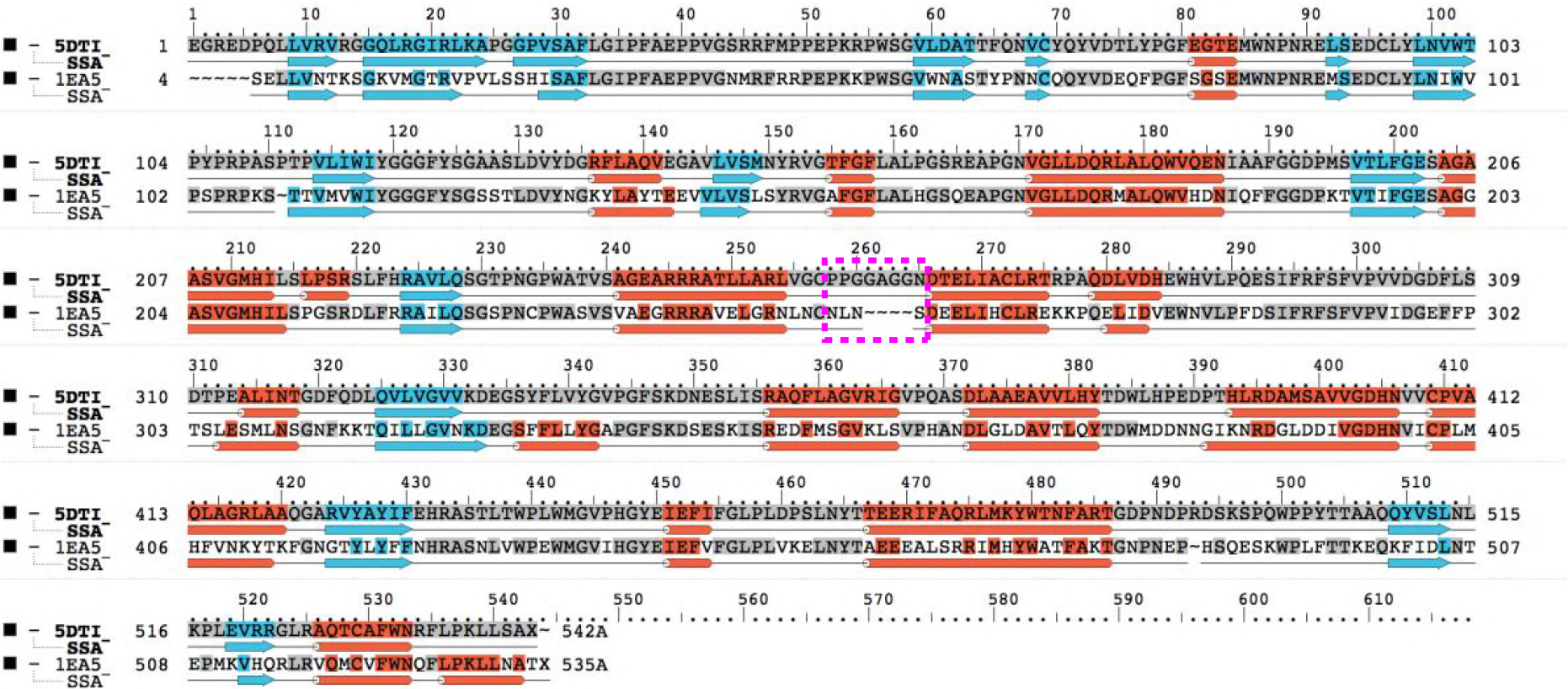
Pairwise structure-based sequence alignment of *Tc*AChE and mAChE employing the Schrödinger Maestro multiple sequence viewer (MSV) [19]. They possess 58.1% identity and 74.1% similarity. The numbers above the alignment correspond to those of the mAChE sequence. The additional loop in mAChE (PPGGAGGN) is highlighted in magenta. The α-helices, β-strands and loops in the structures of *Tc*AChE and mAChE are represented as cylinders (red), arrows (cyan) and lines (black), respectively.

**Fig 5.**
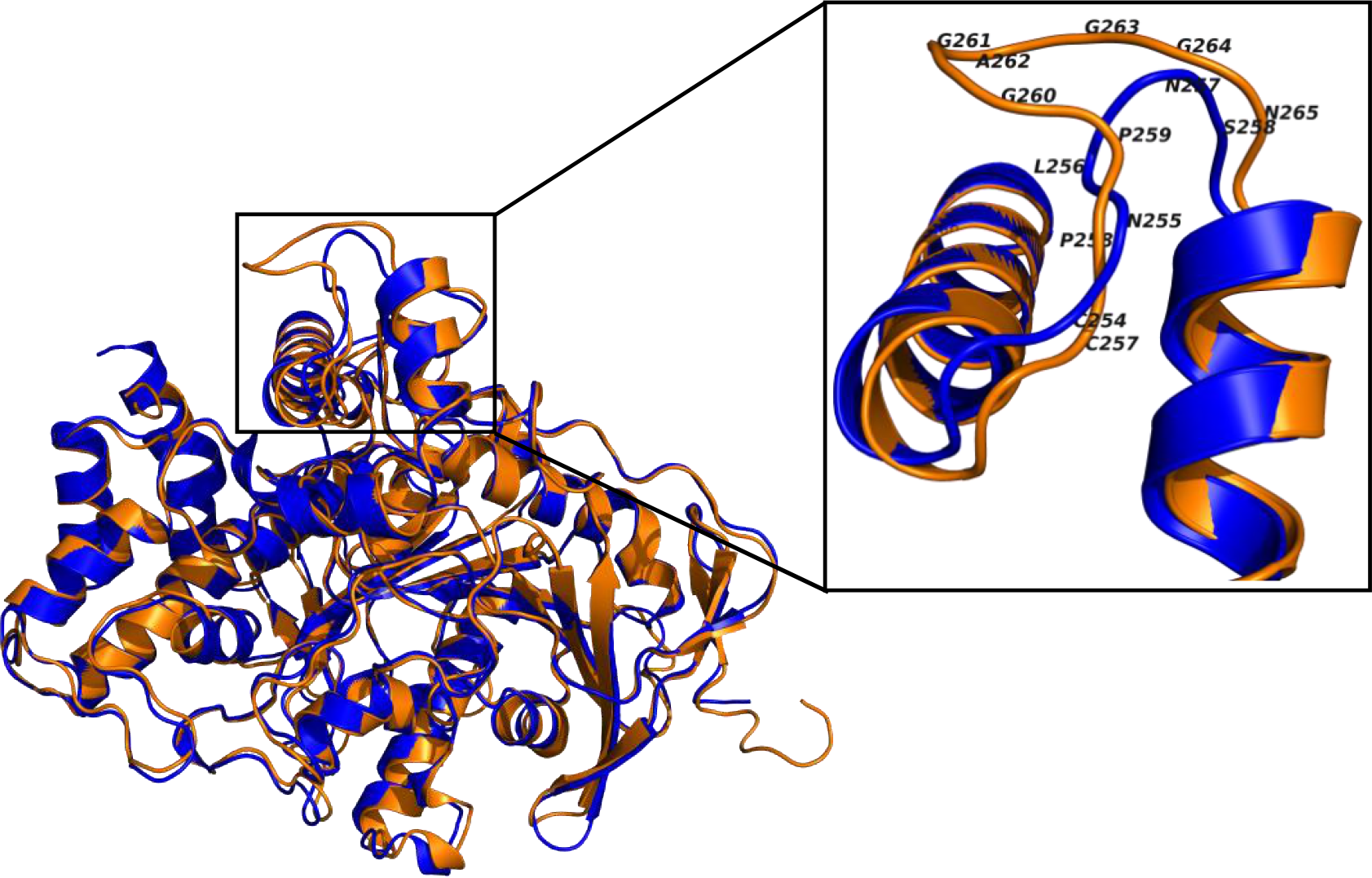
Superposition of cartoon representations of the crystal structures of *Tc*AChE (blue) and mAChE (orange). The inset (upper right) highlights the large loop present in mAChE.

### 2.2. MD simulations of *Tc*AChE and mAChE

Knowledge of the intrinsic flexibility of a protein is important for understanding how structure is linked to function [20]. Accordingly, we performed 20-ns MD simulations on the native crystal structures of *Tc*AChE and mAChE (Fig 6). Although the 3D structures are almost identical, the MD simulations clearly reveal that mAChE is significantly more flexible than *Tc*AChE. Fig 6 shows that the average RMSD for the backbone atoms in mAChE is ~1.3 Å, compared to ~1.0 Å in *Tc*AChE (see S2 Fig in Supplementary Information). Almost throughout the entire sequence the mAChE residues are more flexible than those of *Tc*AChE (Fig 7) (see S3 Fig in Supplementary Information). One of the most flexible regions is around the inserted PPGG sequence in mAChE (Fig 7).

**Fig 6.**
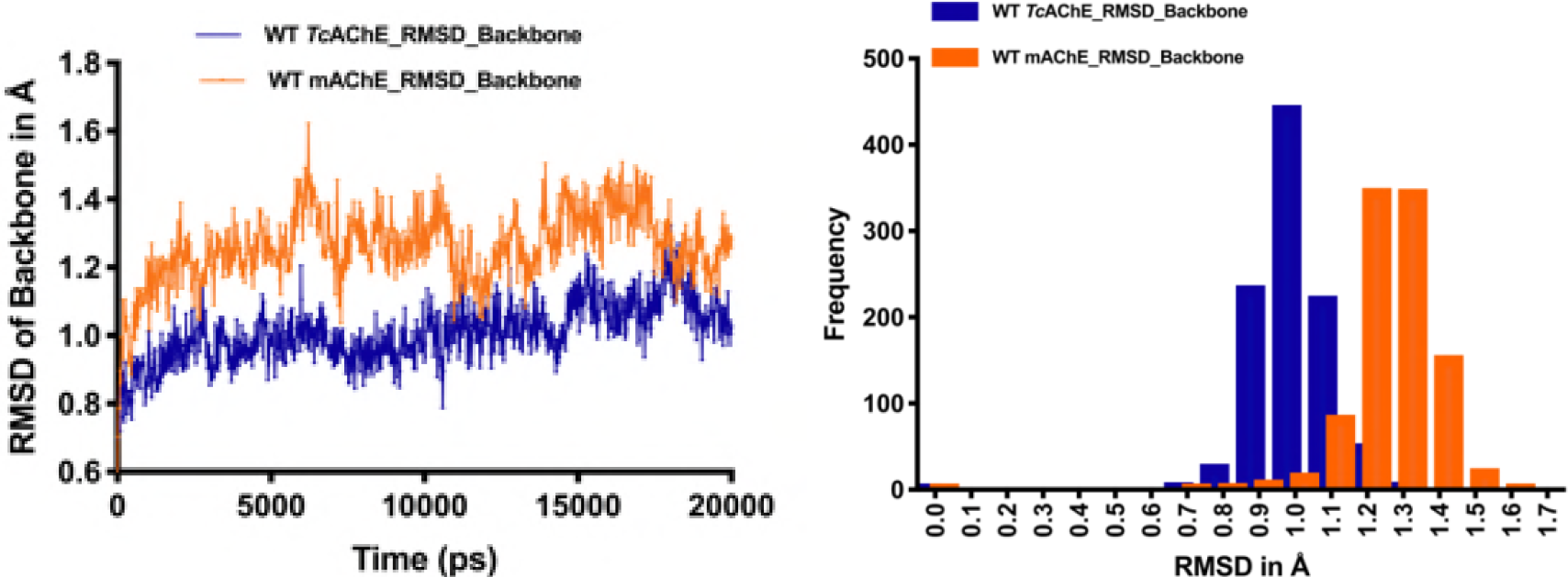
RMSD values for backbone atoms of mAChE (orange) and *Tc*AChE (blue) obtained from the 20-ns MD simulations. Left panel, time course of the simulation; right panel, frequency histograms derived by sampling the data displayed on the left at 20-ps intervals.

**Fig 7.**
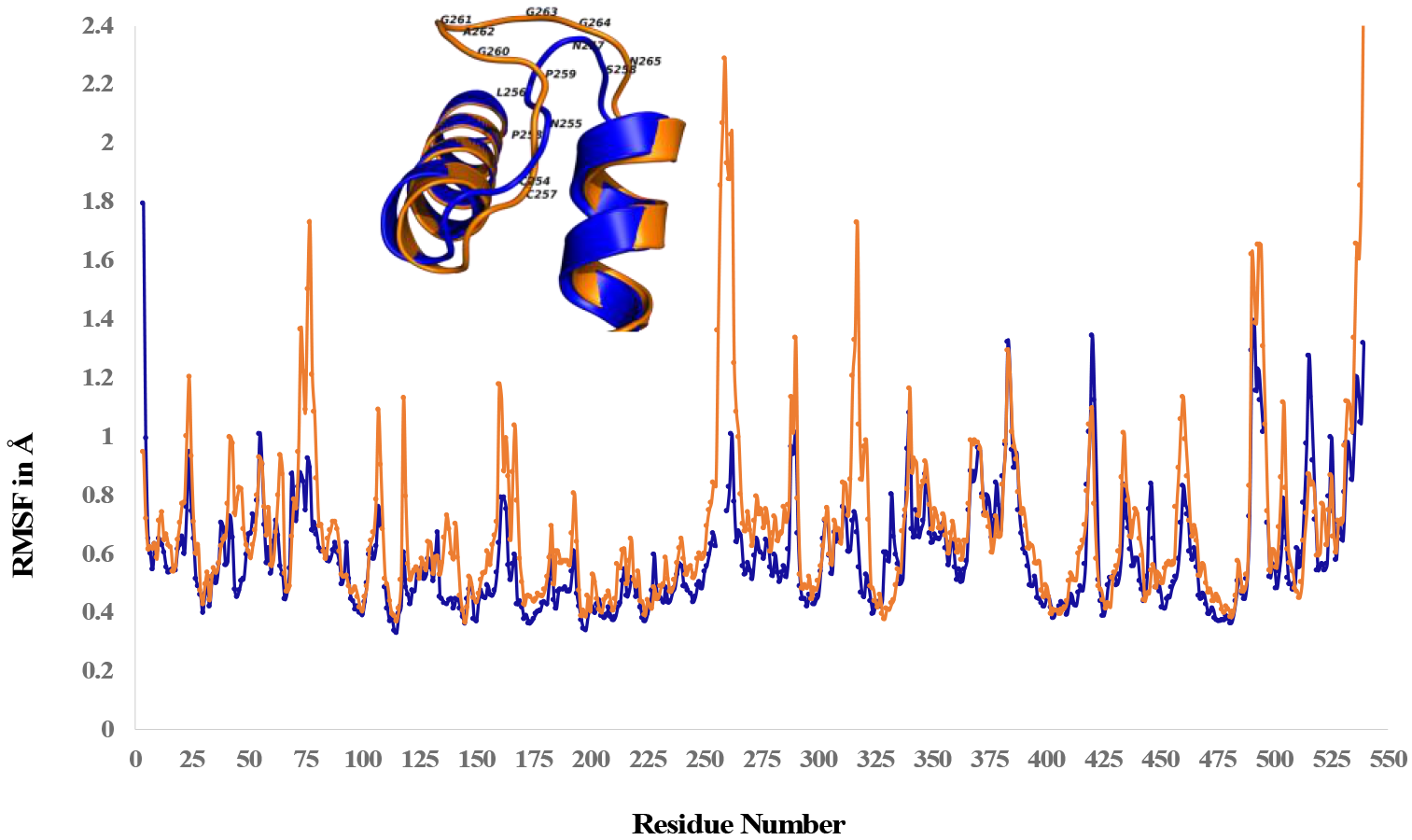
RMSF values for the backbone atoms of mAChE (orange) and *Tc*AChE (blue) obtained from the 20-ns MD simulations.

Our MD simulations are consistent with those previously published for mAChE [21] and for *Tc*AChE [22] (see S4a Fig and S4b Fig in Supplementary Information).

### 2.3. GlideXP docking of PMSF and BSF into *Tc*AChE and mAChE followed by MD simulation

#### 2.3.1. Docking

No crystal structures are available of complexes or conjugates of BSF or PMSF with AChE. Therefore, GlideXP was used to dock both inhibitors into *Tc*AChE [23] and mAChE [24]. In all four cases, the inhibitors dock at a very similar position (Fig 8), with the electrophilic sulfur atom remaining at a great distance from the nucleophilic hydroxyl group of S200(S203) within the active site at the bottom of the gorge (Fig 8 and Table 1).

**Table 1:** Distances between S200(S203)*Oγ* of *Tc*AChE and mAChE and the sulfur atom of BSF or PMSF after docking alone or after docking followed by MD simulation.

|  | *TcAChE* |  | m*AChE* |  |
| --- | --- | --- | --- | --- |
|  | Docked | MD | Docked | MD |
| BSF | 11.8 Å | 4.5 Å | 10.2 Å | 3.0 Å |
| PMSF | 11.8 Å | 11.2 Å | 10.2 Å | 4.0 Å |

**Fig 8.**
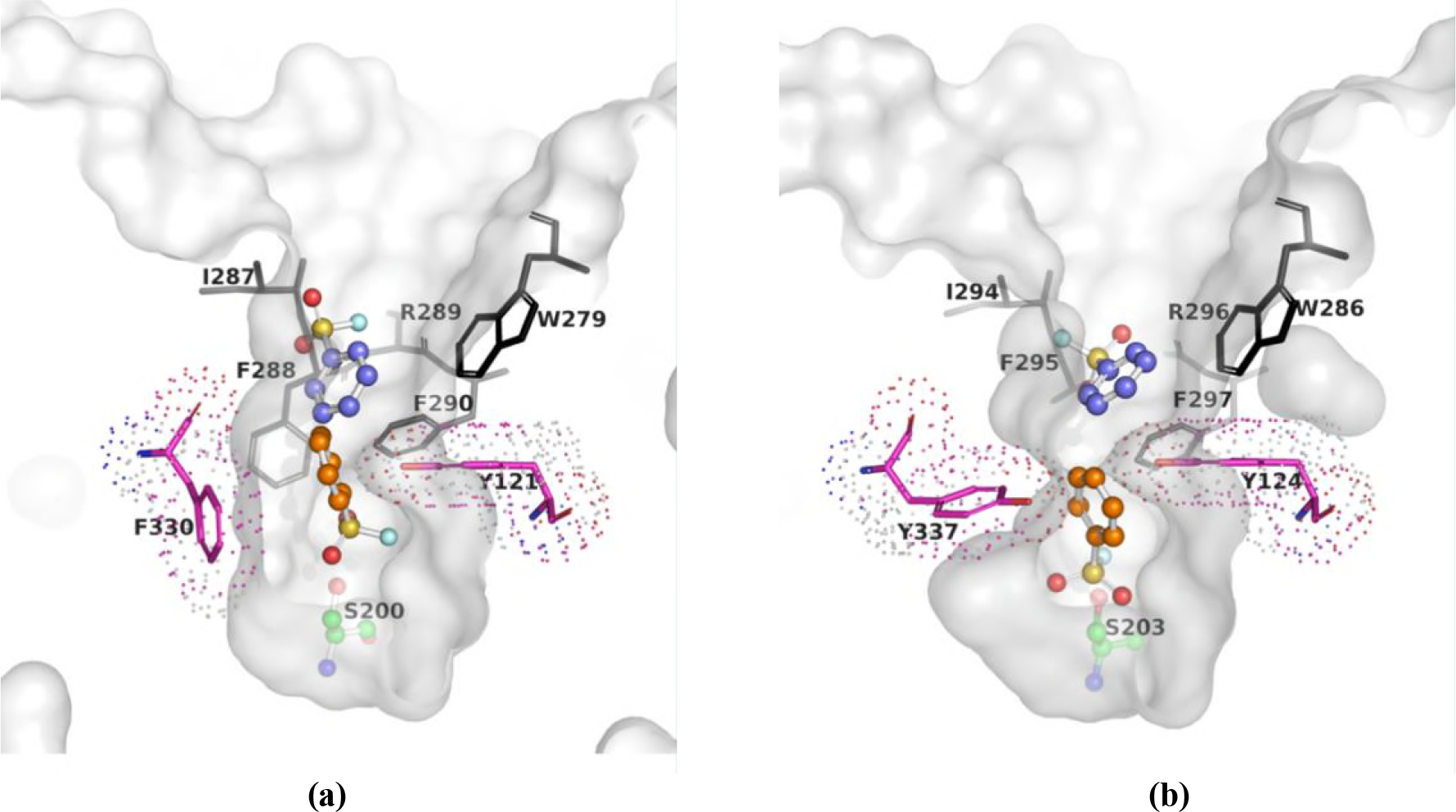
GlideXP docking and MD simulation for interaction of BSF and PMSF with *Tc*AChE and mAChE. The upper panels show the data obtained for interaction of BSF with (a) *Tc*AChE and (b) mAChE. The lower panels show the data obtained for interaction of PMSF with (c) *Tc*AChE and (d) mAChE. In all four panels two copies of the ligand are displayed. One shows the position of the ligand after docking alone (blue), and the other shows the position after docking followed by MD simulation (orange). It should be noted that the orientations of the amino-acid side-chains displayed are those seen prior to the MD simulations.

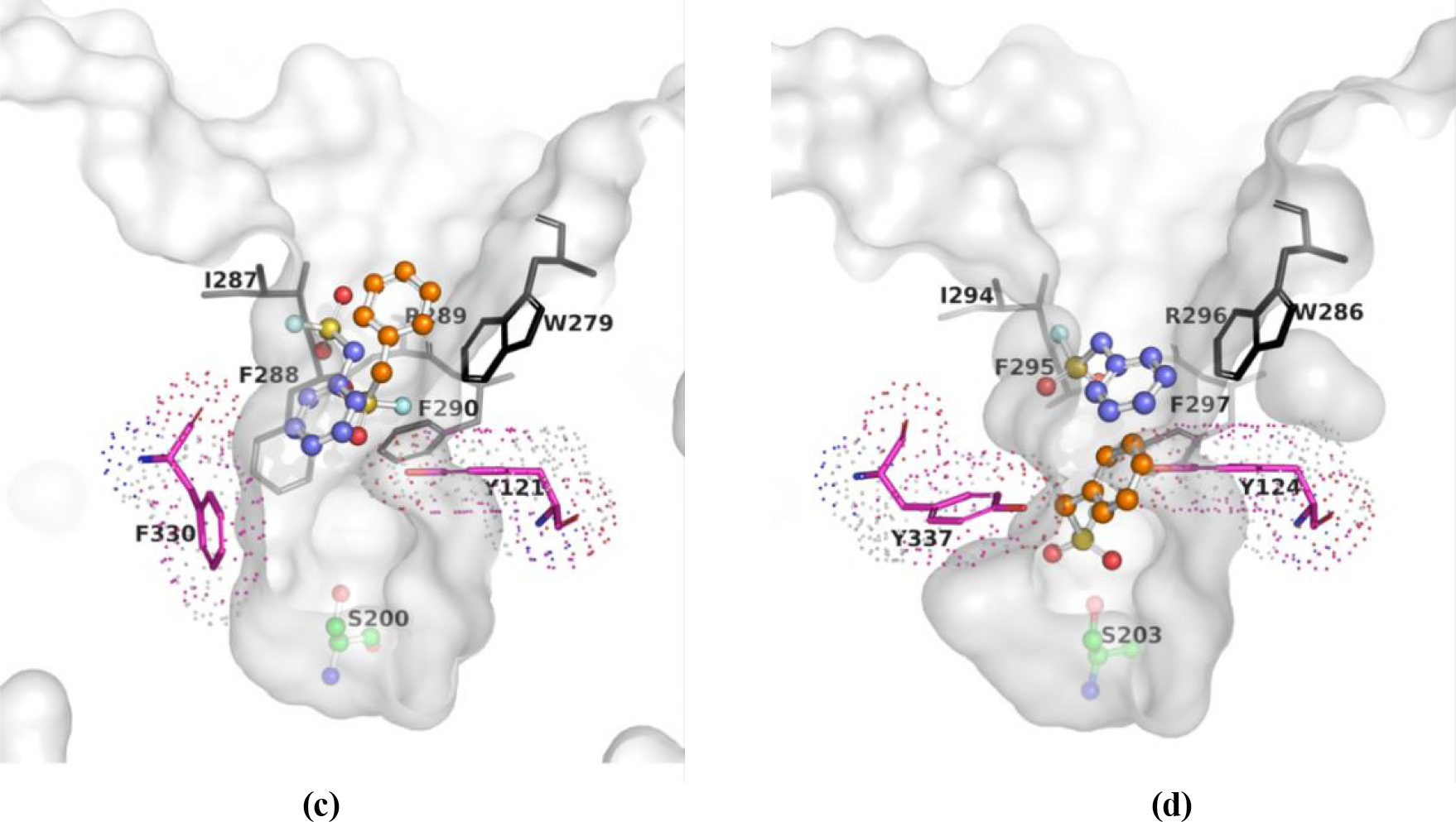

#### 2.3.2. MD Simulation Following Docking

Since BSF inhibits both *Tc*AChE and mAChE, whereas PMSF inhibits only mAChE, the docking results displayed in the previous section obviously do not reflect the experimental data. We decided, therefore, to perform MD simulations following docking [24, 25]. It can be clearly seen that after the MD simulation BSF approaches close to S200(S203)*O*γ in both *Tc*AChE and mAChE. However, PMSF approaches S203*O*γ in mAChE, whereas it remains distant from S200*O*γ in *Tc*AChE (Fig 8) (see S5 Fig in Supplementary Information).

The MD values displayed refer to the closest approach of the sulfur atom of the ligand to the active-site S200(S203)*O*γ throughout the entire MD trajectory.

Figs 9 and 10 displays the MD trajectories for the four cases analyzed, and the histograms derived from them. It can be seen that in the interaction of BSF with *Tc*AChE the ligand approaches S200*O*γ quite closely for part of the time, whereas PMSF remains much further away, above the bottleneck. In the corresponding histograms for interactions with mAChE, the trajectory for BSF almost merges with the magenta line corresponding to the VDW contact distance, while that for PMSF is a little further away. In all four cases, after docking alone the sulfonyl group points up the gorge, away from the active-site, whereas after docking it points towards the active-site.

**Fig 9.**
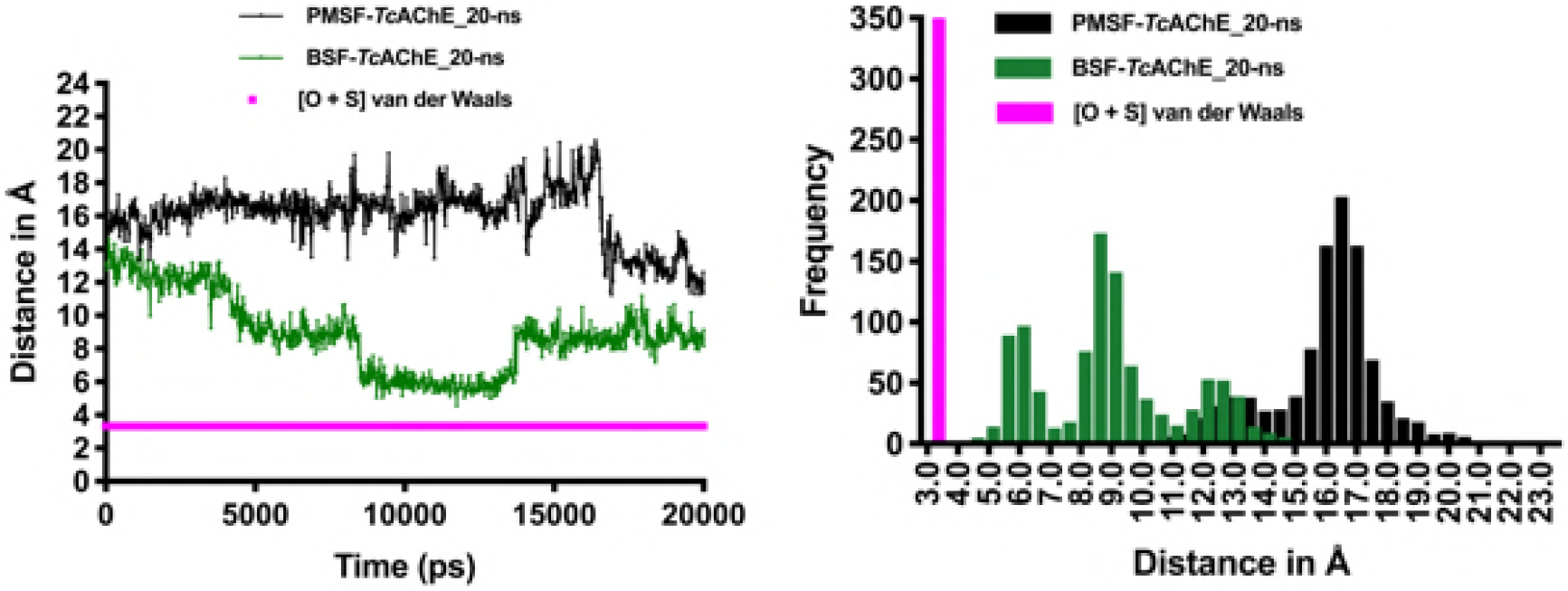
MD trajectories for interaction of PMSF and BSF with *Tc*AChE. Left, 20-ns trajectories showing BSF*-Tc*AChE (green) and PMSF*-Tc*AChE (black); the magenta line represents the minimum distance between the sulfur atom of the inhibitor and S200*O*γ in a productive encounter. Right, frequency histograms derived by sampling the data displayed on the left at 20-ps intervals.

**Fig 10.**
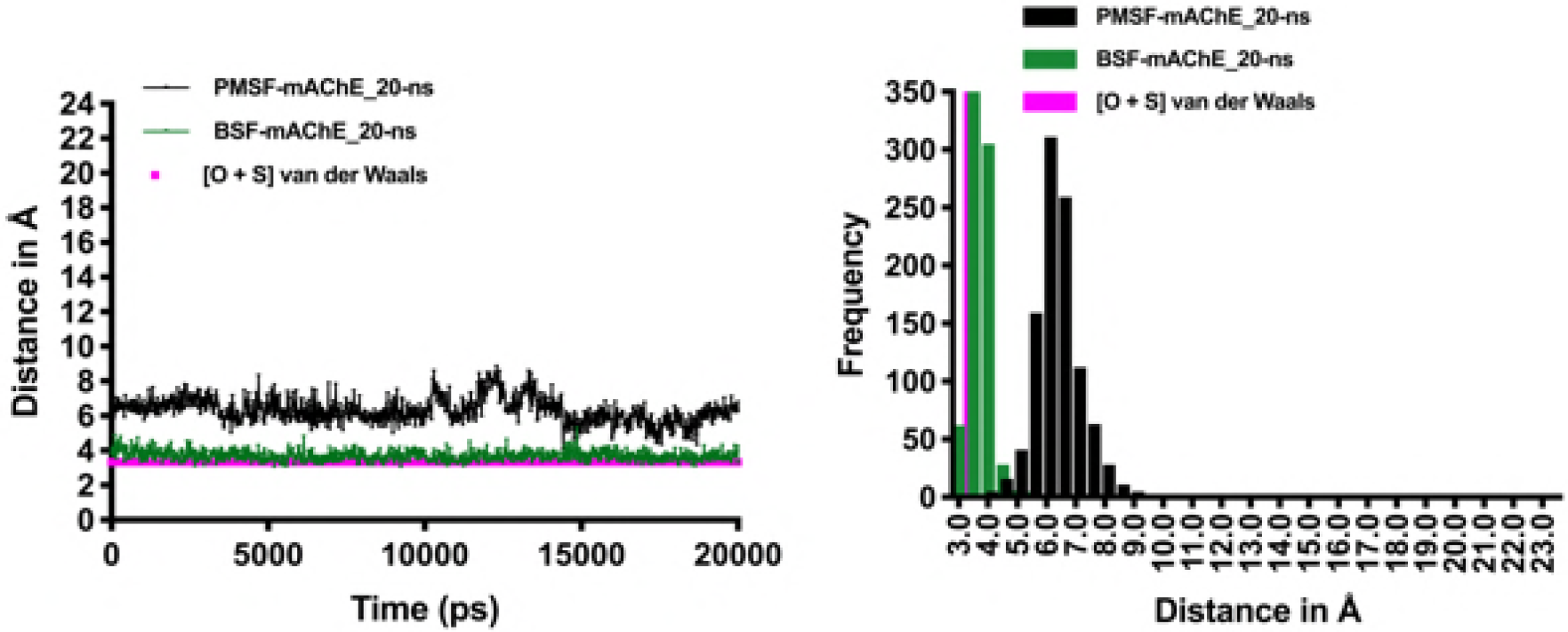
MD trajectories for interaction of PMSF and BSF with mAChE. Left, 20-ns trajectories showing BSF-mAChE (green) and PMSF-mAChE (black); the magenta line represents the minimum distance between the sulfur atom of the inhibitor and S203Oγ in a productive encounter. Right, frequency histograms derived by sampling the data displayed on the left at 20-ps intervals.

## 3. Discussion

As mentioned in the Introduction, *Tc*AChE, mAChE, and human (h) AChE, act at similar rates on their natural substrate, ACh, and on its homolog, ATCh. They have high sequence homology, and their crystal structures reveal almost identical folds. Yet at least two covalent inhibitors interact several orders of magnitude more rapidly with the mammalian enzymes than with *Tc*AChE [13, 17]. It was on this issue that our theoretical study focused. Since *Tc*AChE and the mammalian enzymes do not differ appreciably in their 3D structures [26], steric factors do not seem to explain these very large differences. Consequently, we considered the possibility that differences in their dynamics might provide an explanation. However, it should be kept in mind that, whereas BSF has a planar geometry (Scheme 1), PMSF has a non-planar geometry due to the presence of the additional methylene group, so that its movement through the bottleneck might be expected to be more restricted than that of BSF.

Figs 6 and 7 clearly show large differences in flexibility between *Tc*AChE and mAChE. mAChE is much more flexible than *Tc*AChE along most of the polypeptide chain, displaying an RMSD for backbone atoms of 1.3 Å *vs* 1.0 Å for *Tc*AChE. Particularly high flexibility is seen for a loop that includes residues P258-P259-G260-G261, which are absent in *Tc*AChE (Figs 6 & 7). We thus considered it plausible that this increased flexibility of the mAChE might permit movement of the PMSF through the bottleneck that would not occur in *Tc*AChE.

The results of docking and MD simulations of the WT enzymes by BSF and PMSF mimic the experimental data; thus, BSF comes close to the active-site serine *O*γ atom in both AChEs, but while this is also the case for PMSF binding to mAChE, for *Tc*AChE it remains above the bottleneck, near the top of the gorge, close to the position it occupied in the docking protocol alone.

It should be noted that in all cases, after docking alone both PMSF and BSF were oriented with their sulfonyl moiety pointing up the gorge, away from the active site. But when docking was followed by the MD protocol, in all cases the molecule flipped ~180°, so that the sulfonyl moiety pointed towards the active-site S200(S203)*O*γ. This was true also for PMSF interacting with *Tc*AChE, even though in this case the ligand did not cross the bottleneck. In the crystal structure of S203A mAChE referred to in the Introduction (PDB ID 2HA4) [6] two copies of the substrate, ACh, are seen, one below the bottleneck, and one trapped above it. That trapped above it is oriented, like PMSF and BSF after docking followed by MD simulation, with its leaving group facing into the bottleneck, and its quaternary group making a π-cation interaction with W286 in the peripheral anionic site.

Experimental mutagenesis studies that we performed earlier also suggested that breathing motions might be involved in controlling access of PMSF to the active site. The double mutation, F288L/F290V, which enlarges the acyl pocket, thus permitting *Tc*AChE to act on butyrylthiocholine [27], renders it even more susceptible to PMSF than the WT enzyme [13]. Furthermore, the L282A mutation, which has lower thermal stability than the WT enzyme [28], is inactivated by PMSF at a rate similar to that at which it inactivates the F288L/F290V mutant [13, 29]. Thus, in several cases increased flexibility of AChE appears to be associated with the capacity to be inhibited by PMSF.

As mentioned in the Introduction, the carbamylating agent, rivastigmine, inactivates hAChE (which is highly homologous to mAChE, even more so than *Tc*AChE) three orders of magnitude faster than it inactivates *Tc*AChE [17]. However, various organophosphates (OPs) do not display such differences in rates of phosphorylation of *Tc*AChE and hAChE. Thus, the rates of inactivation of *Tc*AChE and hAChE by di*iso*propylphosphorofluoridate (DFP) are quite similar [30], and the potent *S* isomers of VX and Russian VX actually inactivate *Tc*AChE a few-fold faster than hAChE (Y. Ashani, personal communication). Another interesting case for comparison is that of (-)-huperzine A (HupA). This is a bulky alkaloid with a rigid structure and diameter of 9.8 Å. Although HupA is a reversible inhibitor, it inhibits AChE with extremely slow rates of association and disassociation [31]. MD and steered MD simulations show that sizeable distortions of the residues along the active-site gorge are required for it to pass the bottleneck [32–34]. Yet the rates of inhibition of *Tc*AChE and hAChE by HupA are very similar [31]. It is obvious that our understanding of how protein function is coupled to protein dynamics is inadequate, to say the least.

## 4. Methods

### 4.1. Docking Simulations

The crystal structures of *Torpedo californica* acetylcholinesterase (*Tc*AChE, PDB ID: 1EA5 [35, 22], 1.8 Å resolution) and mouse acetylcholinesterase (mAChE, PDB ID: 5DTI [36], 2.0 Å resolution) were retrieved from the RCSB Protein Data Bank (http://www.rcsb.org). The monomers of Chain A were used for all the simulations [37].

#### Protein Preparation for Docking and Molecular Dynamics

The 1EA5 and 5DTI structures were prepared with Protein Preparation Wizard [38, 39] in Maestro-v11.1 [40] prior to grid-based-ligand docking (GLIDE) [41, 42] and MD studies, with the following modifications:

- All waters and cofactors were removed.
- Assignment of bond orders
- Addition of hydrogen atoms
- Indicating the 3 intramolecular disulfide bonds
- Adding unseen atoms of side chains in both *Tc*AChE and mAChE
- Addition of residues 259-264 (PGGAGG) which are not seen in any mAChE crystal structure
- The N and C-termini were capped with acetyl (ACE) and N-methyl-amino (NMA) groups, respectively
- Assignment of ionization states
- All-atom restrained molecular mechanics (MM) minimizations was performed with a termination criterion of 0.18 Å on the heavy (non-hydrogen) atoms by employing the OPLS3 force field [43]

#### Ligand preparation

Geometries of the two inhibitors, BSF and PMSF, were optimized using the PBE0 hybrid density functional method [44, 45] combined with the cc-pVTZ Gaussian basis set [46]. Calculations were performed with water as the solvent, using the SCRF continuum solvation model [47]. All calculations utilized the Gaussian 09 suite [48]. The parameters of BSF and PMSF were assigned using the OPLS3 force field [43] with the LigPrep [49] tool available in the Schrödinger suite.

#### Grid Preparation

Due to the large dimensions of the active-site gorge of AChE, we changed the Glide grid box sizes from the default values [50] (see S6 Fig in Supplementary Information) shown in Fig 11a to those shown in Fig 11b, to ensure that all ligand poses would be located within the purple box. “The outer, purple box defines the volume in which the grid potentials are computed. All ligand atoms of a valid pose must be located within this outer box. The inner, green box defines the volume that the ligand center explores during the exhaustive site-point search” (https://www.schrodinger.com/kb/701).

**Fig 11.**
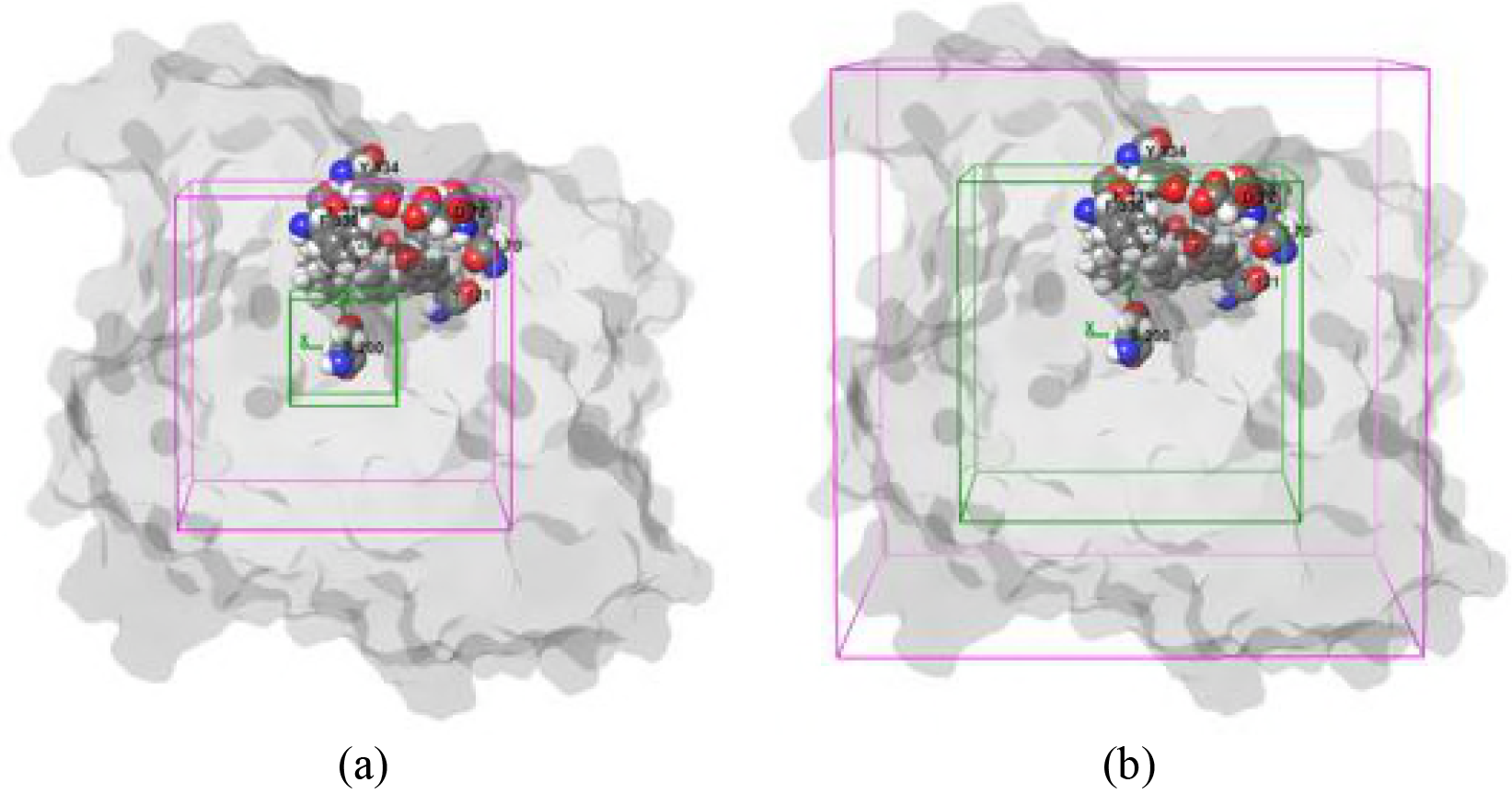
Glide grid box settings employed: The boxes were centered around S200(S203)^1^, the active-site serine at the bottom of the gorge. (a) Default settings of the inner (green) and outer (magenta) grid boxes were 10×10×10Å and 30×30×30Å, respectively; (b) Inner and outer grid box settings employed for the docking studies on *Tc*AChE and mAChE were 30×30×30Å and 50×50×50Å, respectively.

#### Glide Docking

Docking, with a rigid protein structure and a flexible ligand, was performed using Schrödinger Glide-v7.4, employing the XP (extra precision) option [42]. One hundred poses were generated per docking calculation. In order to soften the potential for non-polar atoms of the protein (within the grid points) and the ligands, their VDW radii were scaled to 1.0 Å, with a partial charge cut-off of 0.25 units, while VDW radii of remaining atoms were not scaled (see S7 Fig in Supplementary Information). Glide docking uses a hierarchical clustering algorithm to produce the best set of ligand-binding locations in the defined receptor grid space. The lowest XP glide score, for a given ligand, indicates the highest binding affinity for the enzyme.

### 4.3. Molecular Dynamics Simulations

The Desmond-v4.9 [51] simulation package from Schrödinger was used for all MD simulations. The protein-ligand complex systems were solvated using the explicit SPC [52] solvent model in a 10×10×10Å box, employing the OPLS3 force field [43] (see S8 Fig in Supplementary Information). The charge of the system was neutralized by adding 0.15 M NaCl. The Desmond standard NPT relaxation protocol was used, except that the equilibration time was extended from 24 ps to 5 ns [53] (see S9 Fig in Supplementary Information). After the equilibration step, 20 ns MD simulations were performed (see S10 Fig in Supplementary Information). The simulations were run on the Faculty of Chemistry HPC (High-Performance Computing) Facility at the Weizmann Institute of Science.

## Supporting Information

**S1 Fig.** Cartesian coordinates, in Å, of the two inhibitors, BSF and PMSF, were optimized using the PBE0 hybrid density functional method combined with the cc-pVTZ Gaussian basis set in aqueous phase, as described in the paper.

**S2 Fig.** RMSD values, in Å, for backbone atoms of mAChE and *Tc*AChE obtained from the 20-ns MD simulations (see Fig 6 in paper).

**S3 Fig.** RMSF values, in Å, for the backbone atoms of mAChE and *Tc*AChE obtained from the 20-ns MD simulations (see Fig 7 in the paper).

**S4a Fig.** (a) RMSD of all heavy atoms (blue) and Cα atoms (orange) from the native mAChE structure over a 20-ns MD simulation with OPLS3 force field at 20-ps time intervals in the trajectory. (b) Time dependence of RMSD deviations of all heavy atoms (A) and Cα atoms (B) from the crystal structure of mAChE [21]. The thick line represents the unliganded simulation, and the thin line represents the liganded (huperzine A) simulation. (c) RMSD of all heavy atoms (blue) and Cα atoms (orange) from the native structure for the first ns of the traces shown in (a).

**S4b Fig.** (a) RMSF of Cα atoms of unliganded *Tc*AChE in a 20-ns MD simulation using the OPLS3 force field, Nose-Hoover chain thermostat and Martyna-Tobias-Klein barostat, as in the body of the text; (b) RMSF of Cα atoms of unliganded *Tc*AChE [22] in a 20-ns MD simulation using the GROMOS96 force field, Berendsen thermostat and Berendsen barostat.

**S5 Fig.** Distances, in Å, between S200(S203)*Oγ* of *Tc*AChE and mAChE and the sulfur atom of BSF or PMSF after docking followed by MD simulation.

**S6 Fig.** Grid box parameters in the GLIDE docking program.

**S7 Fig.** The ligand docking parameters in the GLIDE XP docking program.

**S8 Fig.** The parameters for adding solvent to the ligand-protein complexes.

**S9 Fig.** Protocol and parameters of NPT relaxation after adding solvents to the docked ligand-protein complex.

**S10 Fig.** Protocol and parameters of NPT Production MD simulations after NPT relaxation processes

## Acknowledgments

JMLM acknowledges funding from the Israel Science Foundation (grant 1358/15) and from the Weizmann Institute’s SABRA (Supporting Advanced Basic Research) program, the latter supported by a grant from the Estate of Emile Mimran. The funders had no role in study design, data collection and analysis, decision to publish, or preparation of the manuscript. We thank Prof. Koby Levy and Dr. Chakrapani Subramanyam for valuable discussions.

## Author Contributions

Conceptualization: Israel Silman, Joel L. Sussman, Nellore Bhanu Chandar. Formal analysis: Nellore Bhanu Chandar, Irena Efremenko.
Funding acquisition: Jan M.L. Martin.
Investigation: Nellore Bhanu Chandar, Irena Efremenko, Israel Silman, Jan M.L. Martin, Joel L. Sussman.
Methodology: Nellore Bhanu Chandar, Irena Efremenko. Project administration: Jan M.L. Martin.
Supervision: Israel Silman, Jan M.L. Martin, Joel L. Sussman. Validation: Israel Silman, Jan M.L. Martin, Joel L. Sussman. Visualization: Joel L. Sussman.
Writing - original draft: Nellore Bhanu Chandar, Joel L. Sussman, Israel Silman.
Writing - review & editing: Nellore Bhanu Chandar, Joel L. Sussman, Israel Silman, Jan
M.L. Martin.

## Footnotes

1 Throughout the text, when reference is made to amino acid residues in both *Tc*AChE and mAChE, the *Tc*AChE residue appears first, followed by the corresponding mAChE residue in brackets, *e.g.*, S200(S203).

